# Generating publication ready visualizations for Single Cell transcriptomics using SCpubr

**DOI:** 10.1101/2022.02.28.482303

**Authors:** Enrique Blanco-Carmona

## Abstract

Single Cell transcriptomic analysis has become a widespread technology of choice when it comes to understanding the differences at a transcriptomic level in heterogeneous samples. As a consequence, a plethora of analysis tools have been published to tackle the different analysis steps from count matrix generation to downstream analysis. Many of them provide ways to generate visualizations of the data. While some design choices are made, it is a common practice to provide the user with visualizations as raw as possible so that they can be customized to the user needs. However, in many cases these final customization steps are either time consuming or demand a very specific set of skills. This problem is addressed by *SCpubr*, which sacrifices some of this initial freedom of choice in aesthetics to provide the user a more streamlined way of generating high quality Single Cell transcriptomic visualizations.

## Introduction

The field of Single Cell transcriptomics is currently evolving at an incredible speed. Over the course of several years, a wide range of analysis tools has emerged and consolidated as reference tools for general use. Examples of this are scanpy [1] for python and Seurat [2] for R. While this field of research is in constant improvement, a set of key visualizations has been established as standard and is applied transversely across any analysis performed on Single Cell data sets. Examples of this include the ability of visualizing the cells in a dimensional reduced embedding such as UMAP [3] either as categorical or continuous variables, displaying the expression profile of genes as violin plots or dot plots — if a second variable such as the cells expressing the gene in the group is present — or even querying the cell type composition of a single cell data set using a bar plot.

These types of visualizations are considered basic, and are therefore implemented in many of the available software (packages), even when their end goal is focused on the analysis part rather than the visualization. In this context, a package solely dedicated to improving the aesthetics of these widely used visualizations has yet to be designed or to gain popularity. While aesthetics is mostly a subjective topic, it is a matter of fact that achieving great aesthetics in figures in a programmatic way requires a reasonable investment of time, for which *SCpubr* has been designed. As an R package, it is fully compatible with Seurat objects and produces high quality visualization of the most common visualizations in Single Cell transcriptomics.

## Methods

For this publication, a publicly available data set from 10X Genomics [4] containing 10000 human peripheral blood mononuclear cells (PBMC) was used. Count matrix was generated using *cellranger v6*.*1* [5] and analyzed using Seurat v4. Cells for which the number of unique molecular identifiers (UMIs) was lower than 1000, the number of genes was lower than 500 and the percentage of mitochondrial RNA was higher than 20% were excluded from the analysis. Gene expression was normalized using Regularized Negative Binomial Regression (RNBR) [6] and underwent dimensional reduction by computing a principal component analysis (PCA) [7, 8]. The first 30 principal components (PC) were used as basis for a second dimensional reduction using UMAP. Clusters were identified using Louvain [9] algorithm. Regarding the package creation, all the code has been developed in R and designed to be used with Seurat objects. While some of the figures are created from scratch, many act as a wrapper of other packages, taking as input the figure object being outputted by the package that is wrapped, and further modifying it using ggplot2 [10], ggpubr [11], patchwork [12], colortools [13], among other packages.

## Results

In this section, each of the different functions provided by the package will be presented as results. For each function, the major changes and improvements will be presented.

### Dimensional reduction plot

Dimensional reduction plots are one of the signature features of Single Cell transcriptomic analyses. It allows to represent the cells in a two-dimensional reduction, commonly being UMAP. This is implemented in Seurat by the function *Seurat::DimPlot()*. This function offers a wide range of possible outcomes depending on the user’s input. The major changes implemented in *SCpubr::do_DimPlot()* are at the aesthetic level. Apart from providing a custom color map, axes have been removed in the case of using UMAP reductions, as they always represent the first UMAP component in the X axis and the second on the Y axis. Cells are shuffled by default and a bigger dot size has been implemented (Fig. 1, A-B). Another relevant change is when trying to split the resulting figure into multiple panels according to a group variable. In Seurat, this results in the cells belonging to each value in the group being separated into different panels, but the overall UMAP silhouette of all the cells is lost. This, in certain cases, can be critical information. In *SCpubr::do_DimPlot()*, the silhouette remains as grey-colored cells (Fig. 1, C-D).

**Figure 1.**
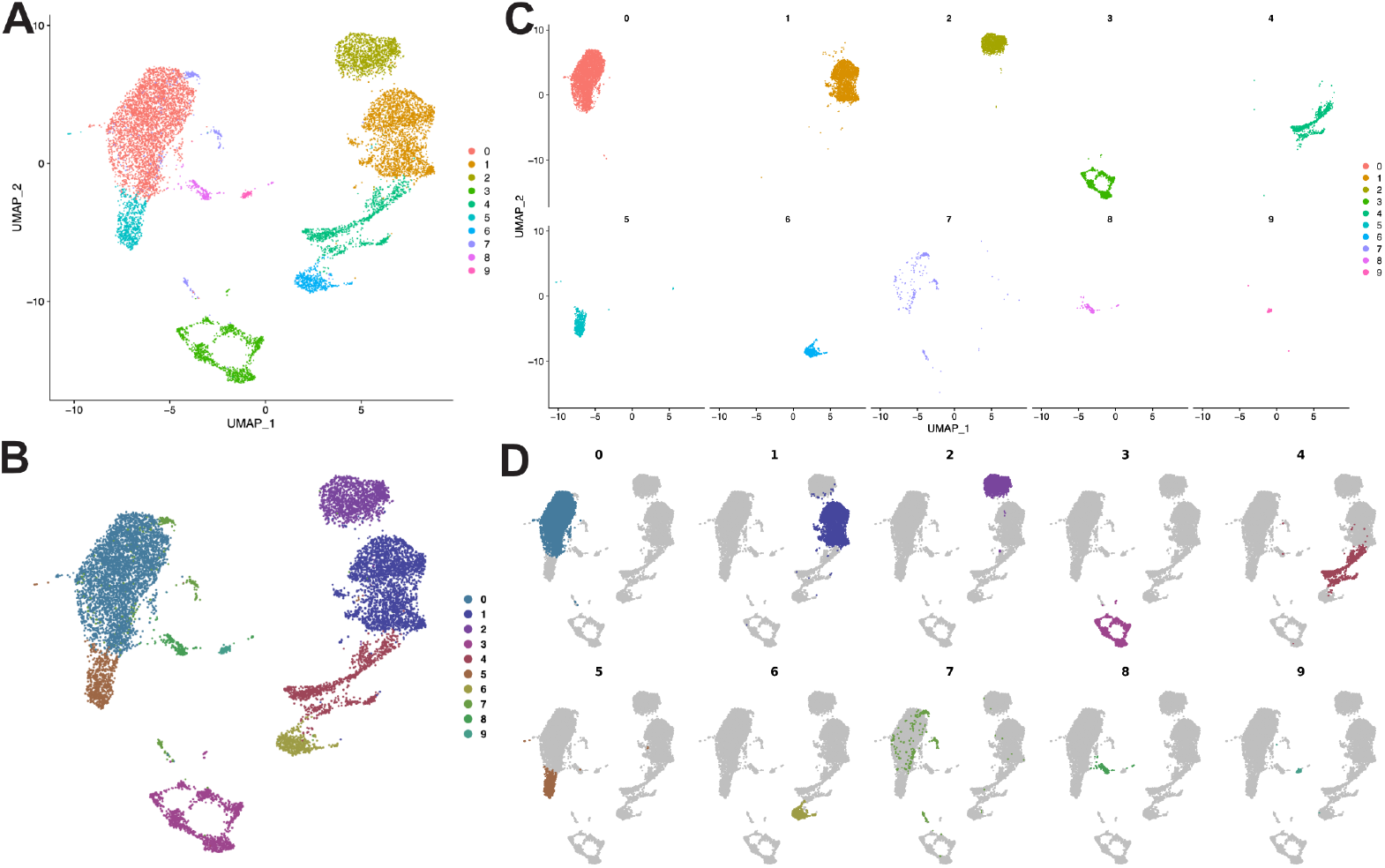
Dimensional reduction plots. **A**. Standard output from *Seurat::DimPlot()*. **B**. Standard output from *SCpubr::do_DimPlot()*. Color map has been modified, axes are removed, dots are bigger in size by default, cells are shuffled by default to avoid identities being plotted in order. **C**. Standard output of *Seurat::DimPlot()* splitting the plot by another variable. **D**. Standard output from *SCpubr::do_DimPlot()* when splitting by another variable. Not selected cells are greyed out and the UMAP silhouette is preserved across panels.

### Feature plots

Feature plots are the counter part of dimensional reduction plots. While the latter focuses on coloring the cells by categorical variables, feature plots color them by continuous variables such as the number of Unique Molecular Identifiers (UMIs), number of genes, expression of a given gene, dimensional reduction coordinates, enrichment scores, etc. This is implemented in Seurat by the function *Seurat::FeaturePlot()*. Same as with *Seurat::DimPlot()*, this function is able to cope with almost any query from the user. From an aesthetic point of view, *SCpubr::do_FeaturePlot()* focuses on removing the axes and making the legend bold for better readability. The color gradient has been substituted by the *viridis* [14] color scale (Fig. 2, A-B). Also, new functionalities are added such as being able to select a subset of cells to plot, while re-scaling the color gradient to contain only the values from the selected cells (Fig. 2, C). While it is also possible in *Seurat::FeaturePlot()* to split the feature plot into different panels according to a variable defining groups, it once again loses the UMAP silhouette. Furthermore, the color scale applied to all panels is the same, therefore making the colors comparable across panels, but the legend showing the range of values is lost. This is addressed in *SCpubr::do_FeaturePlot()*, by greying out the not selected cells while maintaining the same color gradient across panels (Fig. 2, D-E).

**Figure 2.**
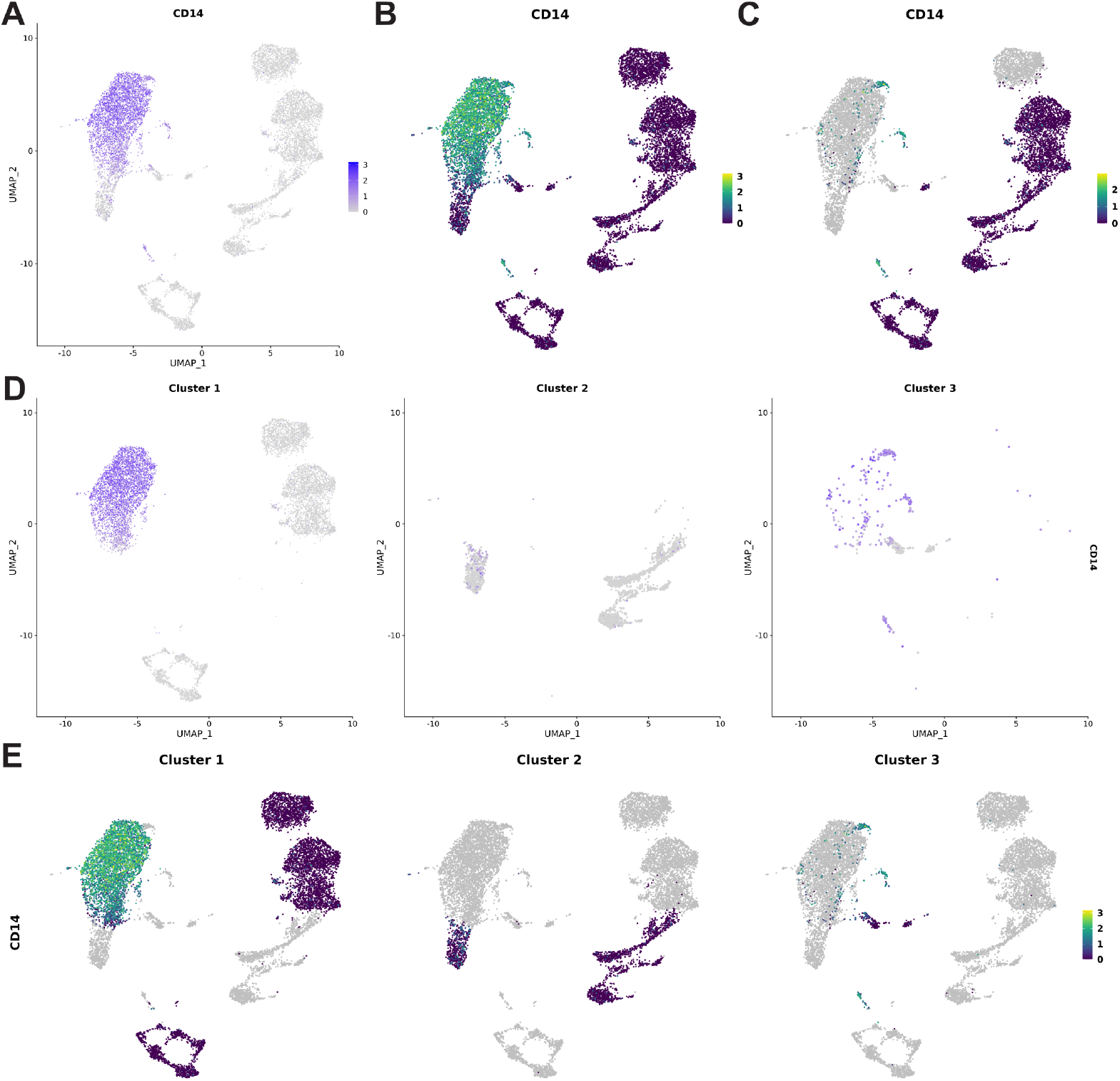
Feature plots. **A**. Standard output from *Seurat::FeaturePlot()*. **B**. Standard output from *SCpubr::do_FeaturePlot()*. Color map has been modified to viridis scale, axes are removed, dots are bigger and legend has bold letters. **C**. Output from *SCpubr::do_FeaturePlot()* when selecting a subset of cells. Cells that are not selected are greyed out and not taken into account when computing the limit of the color scales. **D**. Standard output from *SCpubr::do_FeaturePlot()* when splitting by another variable. UMAP silhouette is lost. **E**. Output from *SCpubr::do_FeaturePlot()* when splitting by another variable. UMAP silhouette is preserved and the legend with the minimum and maximum value is shown.

### Nebulosa plots

Nebulosa plots [15] are a complementary visualization to feature plots. While the latter colors the cells based on a continuous variable, Nebulosa plots computes the density of this variable for each cell. This density value increases depending on the expression of the queried variable in the surrounding cells according to the dimensional reduction. Minor modifications are needed for the output of Nebulosa package, only the removal of the axis and making the legend bold. Most of the value added by *SCpubr::do_NebulosaPlot()* come at either allowing the user to retrieve only the joint density panel or allowing the modification of individual panels’ title (Fig. 3, A-B).

**Figure 3.**
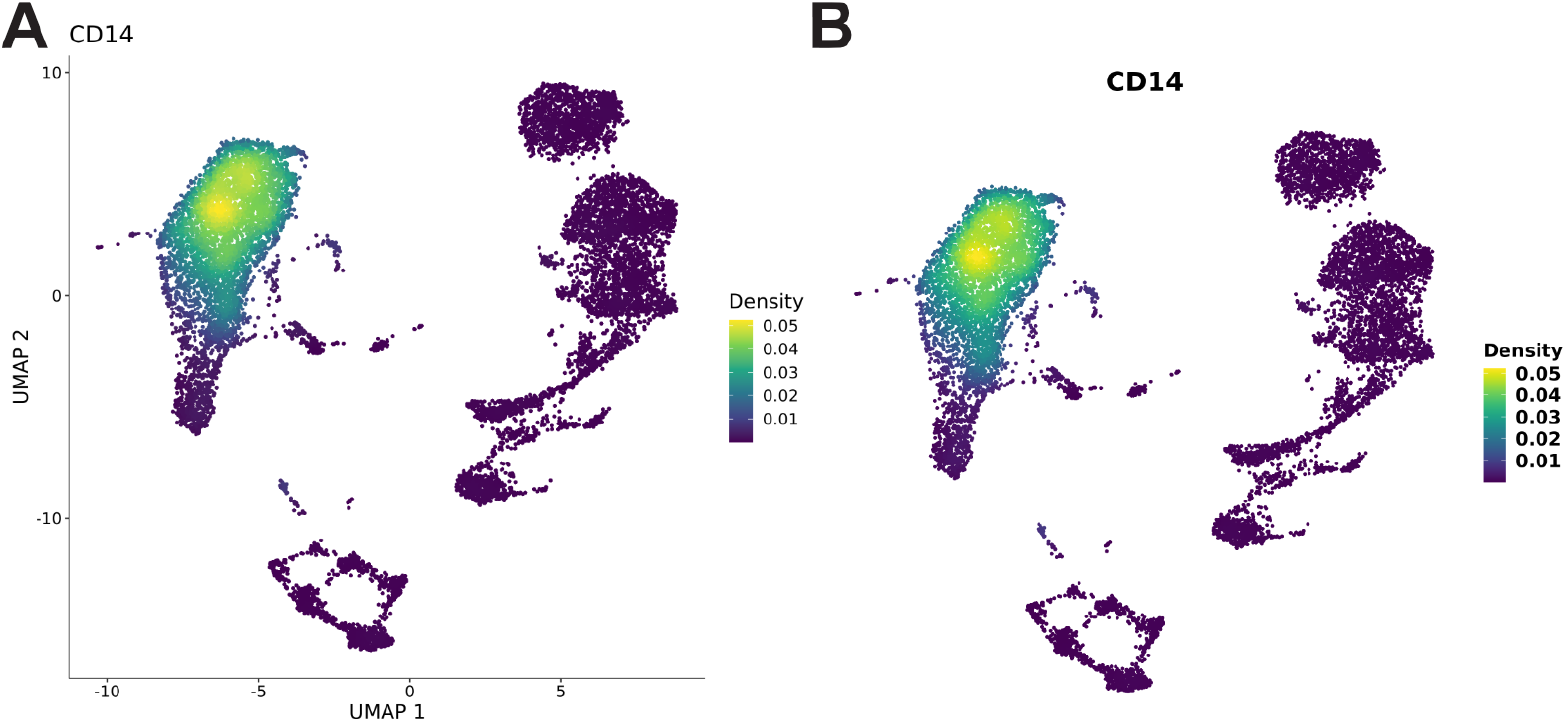
Nebulosa plots. **A**. Standard output from *Nebulosa::plot_density()*. **B**. Standard output from *SCpubr::do_NebulosaPlot()*. Axes are removed and legend has bold letters.

### Bee swarm plots

Bee swarm plots are a really interesting concept for data visualization. It involves ranking a variable for plotting. This is, for a given variable such as a dimensional reduction coordinate, expression values, or enrichment scores, cells are ranked by giving them an order from lowest to highest value. This is then plotted as a modified scatter plot by the *ggbeeswarm* [16] package, separated according to the groups of interest, in which some modifications are made to avoid overplotting of dots, therefore clearly reflecting the density of cells across the range of the ranks. It allows for easy evaluation of the ordering of the cells in the groups along a given variable. It is a visualization that becomes really handy for variables such as pseudotime [17]. Based on this concept, *SCpubr::do_BeeSwarmPlot()* generates this visualization and lets the user decide how to group the cells and whether to color them categorically based on the groups or based on the continuous variable being ranked (Fig. 4, A-B).

**Figure 4.**
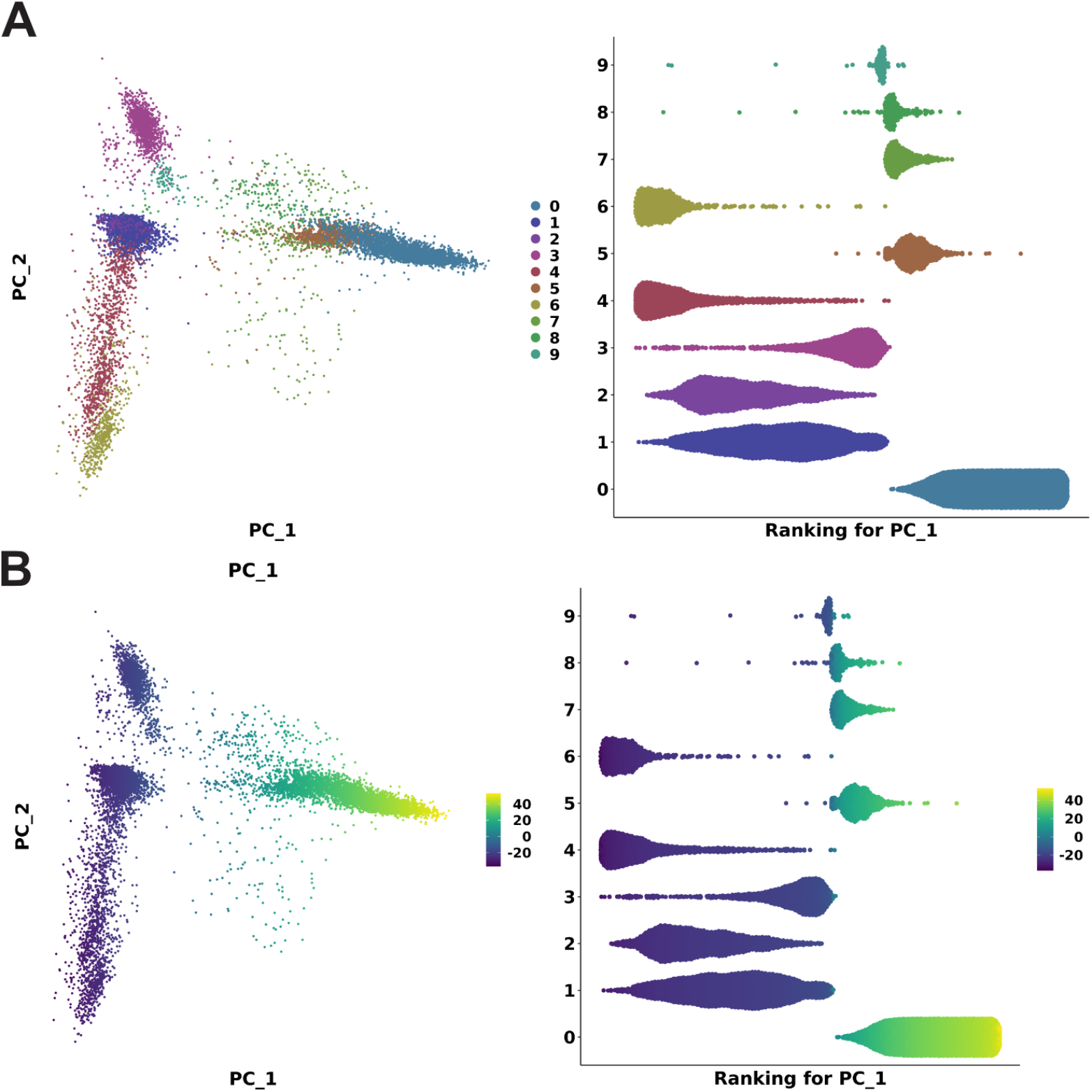
Bee swarm plots. **A**. Side by side comparison of a dimensional reduction plot from *SCpubr::do_DimPlot()* using the PCA dimensional reduction embedding and the output from *SCpubr::do_BeeSwarmPlot()* ranking for PC 1 and coloring by a categorical variable such as the different clusters in the sample. **B**. Side by side comparison of a feature plot from *SCpubr::do_FeaturePlot()* using the PCA dimensional reduction embedding and the output from *SCpubr::do_BeeSwarmPlot()* ranking for PC 1 and coloring by a continuous variable such as the value of the cells for the PC_1.

### Violin plots

Violin plots are another set of widely used data visualizations across research fields. They allow to inspect the distribution of a given variable across different groups of data. In Single Cell transcriptomic analysis, this allows to easily perform quality control (QC) on the different data sets by visualizing the number of UMIs or genes, or querying the expression of given genes across cell identities. A function to represent them is implemented in *Seurat::VlnPlot()*. This function returns a violin plot with dots plotted on top of the violin shapes, leaving it up to the user to decide which size the points should have, if any (Fig. 5, A). In *SCpubr::do_VlnPlot()* the output of *Seurat::VlnPlot()* is used as input and dots are removed by default. Instead, a boxplot is added inside the violin shapes to represent the different quantiles of the data [18]. In addition, it provides the user with an easy way to introduce a horizontal line to indicate cutoffs that were applied during QC (Fig. 5, B).

**Figure 5.**
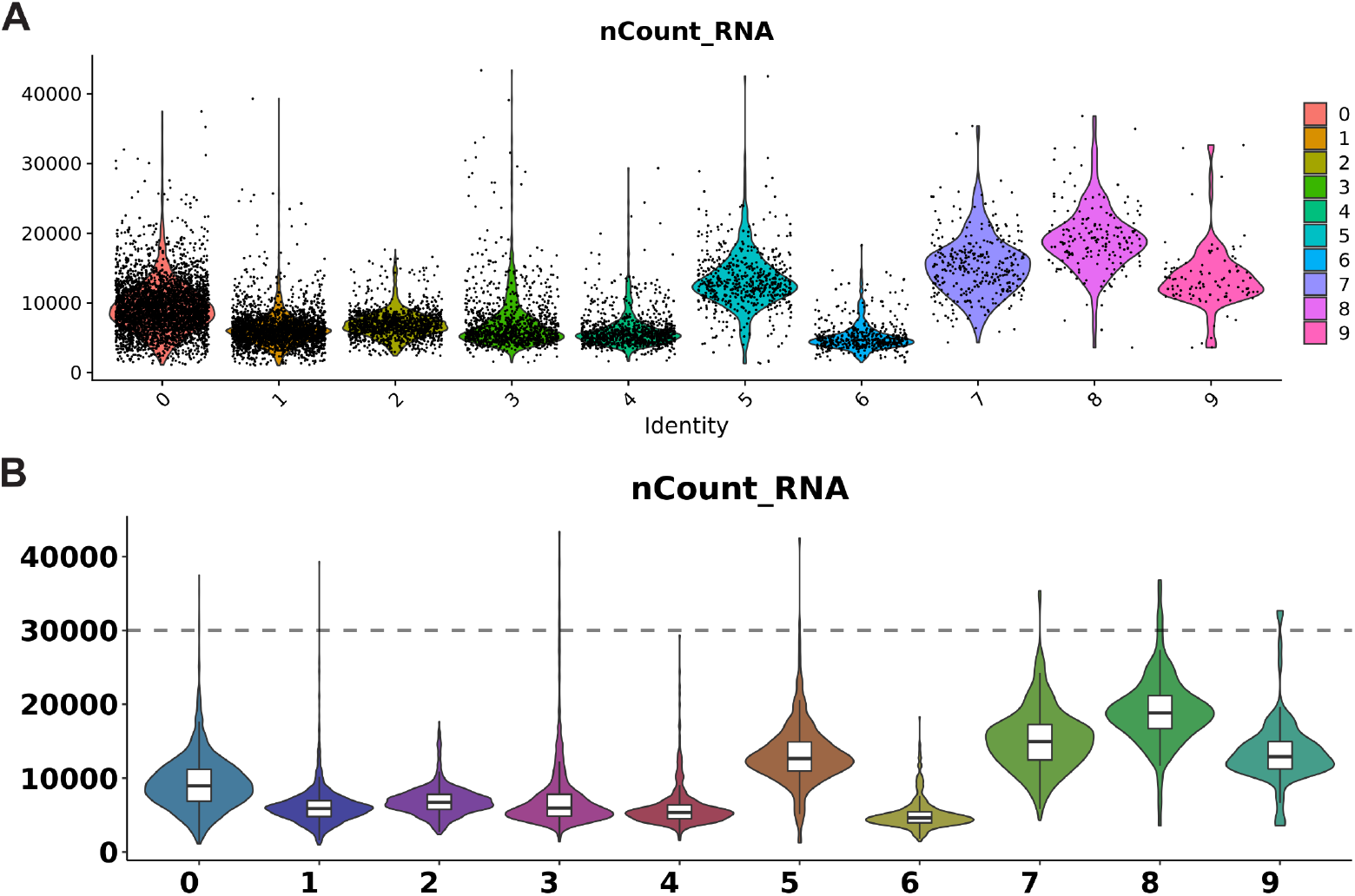
Violin plots. **A**. Output from *Seurat::VlnPlot()* using the number of UMIs as variable to plot. **B**. Output from *SCpubr::do_VlnPlot()* using the number of UMIs as variable to plot and setting a QC cutoff to 30000 UMIs.

### Dot plots

Dot plots are another common visualization, in which values are represented as dots and the size of the dots is mapped to a second variable. The way they are defined in *Seurat::DotPlot()* could be described as a heatmap visualization in which the expression of the genes is displayed in the color scale and instead of cells, we have dots of varying size depending on the percentage of cells in the different groups plotted that express the selected feature. Given the fact that this visualization is really specific for a given purpose, there is not much room for improvement. In *SCpubr::do_DotPlot()*, the output of *Seurat::DotPlot()* is used as input and minor modifications to the color scales, axes and legend are applied (Fig. 6, A-B).

**Figure 6.**
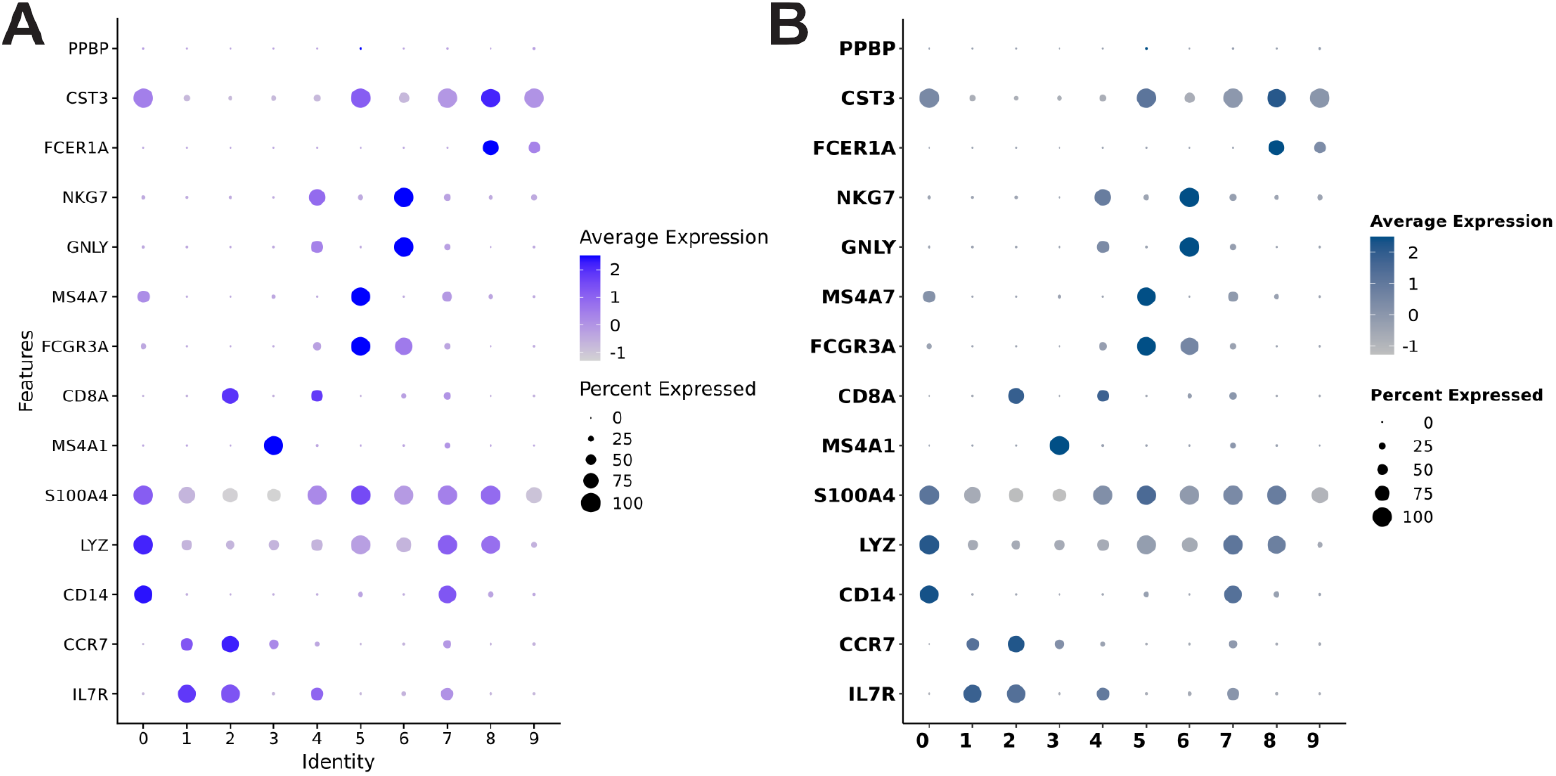
Dot plots. **A**. Output from *Seurat::DotPlot()* using a collection of marker genes and grouping it by each cluster. **B**. Output from *SCpubr::do_DotPlot()* using the same marker genes and groups.

### Bar plots

Bar plots are one of the most basic data visualizations. Yet, they convey a reliable way of displaying numerical data, either in the form of absolute numbers if only one variable is mapped, or also as proportions if a second variable is used to further subdivide each of the bars. In *SCpubr::do_BarPlot()*, simple bars are plotted when a single variable is used and these bars are further divided if a second variable is provided. Furthermore, bars can be shown as absolute numbers or as relative proportions of the groups within each bar (Fig. 7, A-B). As an additional feature, if a second variable is provided, it is also possible to rearrange the bars according to either the descending absolute number or the proportion of a given item of the within-bars groups (Fig. 7, C). Bars can also be displayed either vertically or horizontally.

**Figure 7.**
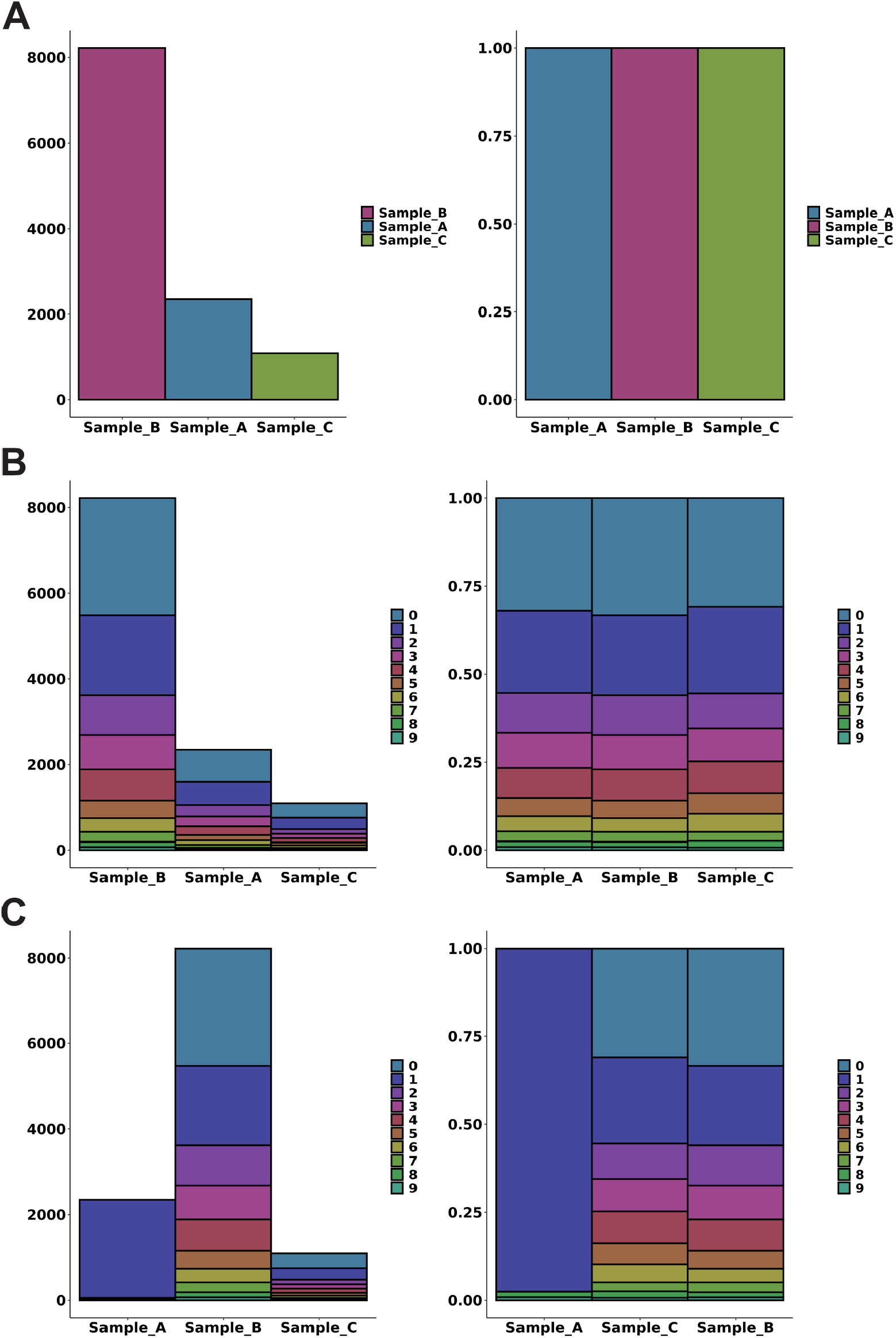
Bar plots. In all panels, arbitrary data sets with different proportions are used. Panel B uses the original inferred clusters while C modifies the proportions. **A**. Output from *SCpubr::do_BarPlot()* using a single variable, in this case the number of cells per data set. **B**. Output from *SCpubr::do_BarPlot()* using two variables. Within each bar, different groups according to the number of cells in each different cluster are drawn. **C**. Output from *SCpubr::do_BarPlot()* using two variables and ordering by the values of cluster 1.

## Conclusions

*SCpubr* is an R package aimed at Single Cell transcriptomic analysis. It uses a Seurat object input for all its functions and returns a high quality visualization. While some aesthetic aspects have been fixed, *SCpubr* offers a wide range of customization features to further tailor the resulting plot to meet the user needs.

## Package availability

*SCpubr* is publicly available for installation in https://github.com/enblacar/SCpubr and a complete reference manual can be found in https://enblacar.github.io/SCpubr-book/. A future release in CRAN is planned.

## Acknowledgements

I thank PD. Dr. med. Pascal Johann, Dr. Natalie Jäger, Dr. Marcel Kool and Prof. Dr. Matthias Schlesner for providing a fruitful working environment that made it possible for this project to take place.

